# Susceptibility of rabbits to SARS-CoV-2

**DOI:** 10.1101/2020.08.27.263988

**Authors:** Anna Z. Mykytyn, Mart M. Lamers, Nisreen M.A. Okba, Tim I. Breugem, Debby Schipper, Petra B. van den Doel, Peter van Run, Geert van Amerongen, Leon de Waal, Marion P.G. Koopmans, Koert J. Stittelaar, Judith M.A. van den Brand, Bart L. Haagmans

## Abstract

Transmission of severe acute respiratory coronavirus-2 (SARS-CoV-2) between livestock and humans is a potential public health concern. We demonstrate the susceptibility of rabbits to SARS-CoV-2, which excrete infectious virus from the nose and throat upon experimental inoculation. Therefore, investigations on the presence of SARS-CoV-2 in farmed rabbits should be considered.

## Text

Severe acute respiratory syndrome coronavirus 2 (SARS-CoV-2) caused a pandemic only months after its discovery in December 2019 (*1*). Slowing down its spread requires a full understanding of transmission routes, including those from humans to animals and vice versa. In experimental settings, non-human primates, ferrets, cats, dogs and hamsters have been found to be susceptible to SARS-CoV-2 infection (*2–4*). Moreover, ferrets, cats and hamsters were able to transmit the virus via the air (*2, 4, 5*). In domestic settings, both dogs and cats have been found to carry the virus, displaying very mild to more severe symptoms, respectively (*5*). Recently, SARS-CoV-2 has been isolated from mink at multiple Dutch farms. Workers at those farms carried viruses that were highly similar to the viruses detected in mink and phylogenetic analyses supported transmission from mink to workers (*6*). Thus, measures to control the spread of SARS-CoV-2 should also include preventing spill over into potential reservoirs, especially since infectious agents can spread rapidly in livestock due to the high densities at which some animals are kept. Given the fact that rabbits are commonly farmed worldwide, we investigated the susceptibility of rabbits to SARS-CoV-2.

## The study

Angiotensin converting enzyme 2 (ACE2) dictates the host range for SARS coronaviruses (*7*), because it engages the viral spike (S) glycoprotein for cell attachment and fusion of the viral and host membranes (*8*). Contact residues of human and rabbit ACE2 critical for binding S are relatively well conserved (*9*). We overexpressed ACE2 from different species on a receptor-deficient cell line followed by SARS-CoV-2 pseudovirus or authentic virus infection. Successful infection of both SARS-CoV-2 pseudovirus and authentic virus was observed for human, rabbit and the Chinese horseshoe bat (*Rhinolophus sinicus*) ACE2 (Figure 1, panel A and B). The ACE2 from a distantly related bat, the great Himalayan leaf-nosed bat (*Hipposideros armiger*), did not support infection and served as a negative control. Transfection of rabbit ACE2 rendered cells susceptible to SARS-CoV-2 infection demonstrated by a clear overlap between infection and ACE2 expression (Figure 1, panel C).

**Figure 1.**
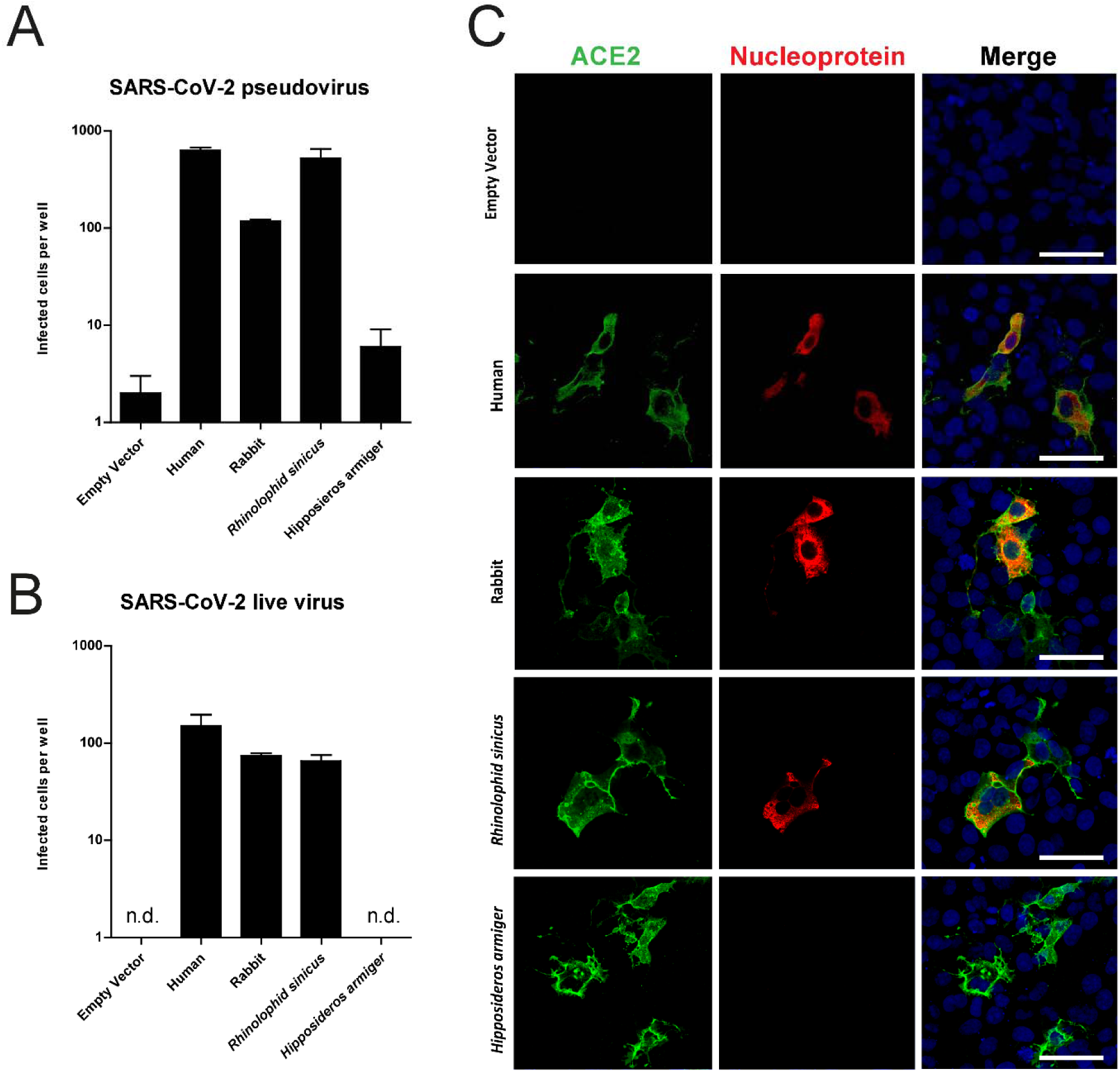
Rabbit ACE2 mediated SARS-CoV-2 infection. SARS-CoV-2 pseudovirus (A) and authen tic virus (B) infection of Cos-7 cells expressing ACE2 of various species. Infectivity was quantified by staining live virus cells with anti-SARS-CoV nucleocapsid and scanning live virus and pseudovirus infected cells. (C) Confocal imaging of ACE2 mediated live virus infection; cells were stained using anti-human ACE2 in green, anti-SARS-CoV nucleocapsid in red and TO-PRO3 in blue to stain nuclei. Scale indicates 50μm.

Next, we inoculated three rabbits with 10^6^ tissue culture infectious dose 50 (TCID_50_) SARS-CoV-2 for a 21 day follow up. None of the inoculated animals showed clinical signs of infection. As shown in Figure 2 (panel A), we found viral RNA in the nose for at least twenty-one days (mean shedding of 15.33 days, SD=5.13), up to fourteen days in the throat (mean shedding of 11.33 days, SD=2.52) and up to nine days in the rectum (mean shedding of 5 days, SD=3.61). Infectious virus shedding from the nose lasted up to seven days (mean shedding of 6.67 days, SD=0.58) with a peak at day two, followed by a second peak at day seven post inoculation (p.i.) (Figure 2, panel B). In the throat, infectious virus was detected only on day one p.i. for one animal. No infectious virus was detected in rectal swabs. All animals followed up to day 21 seroconverted with plaque reduction neutralization test (PRNT_50_) titers of 1:40, 1:320 and 1:640.

**Figure 2.**
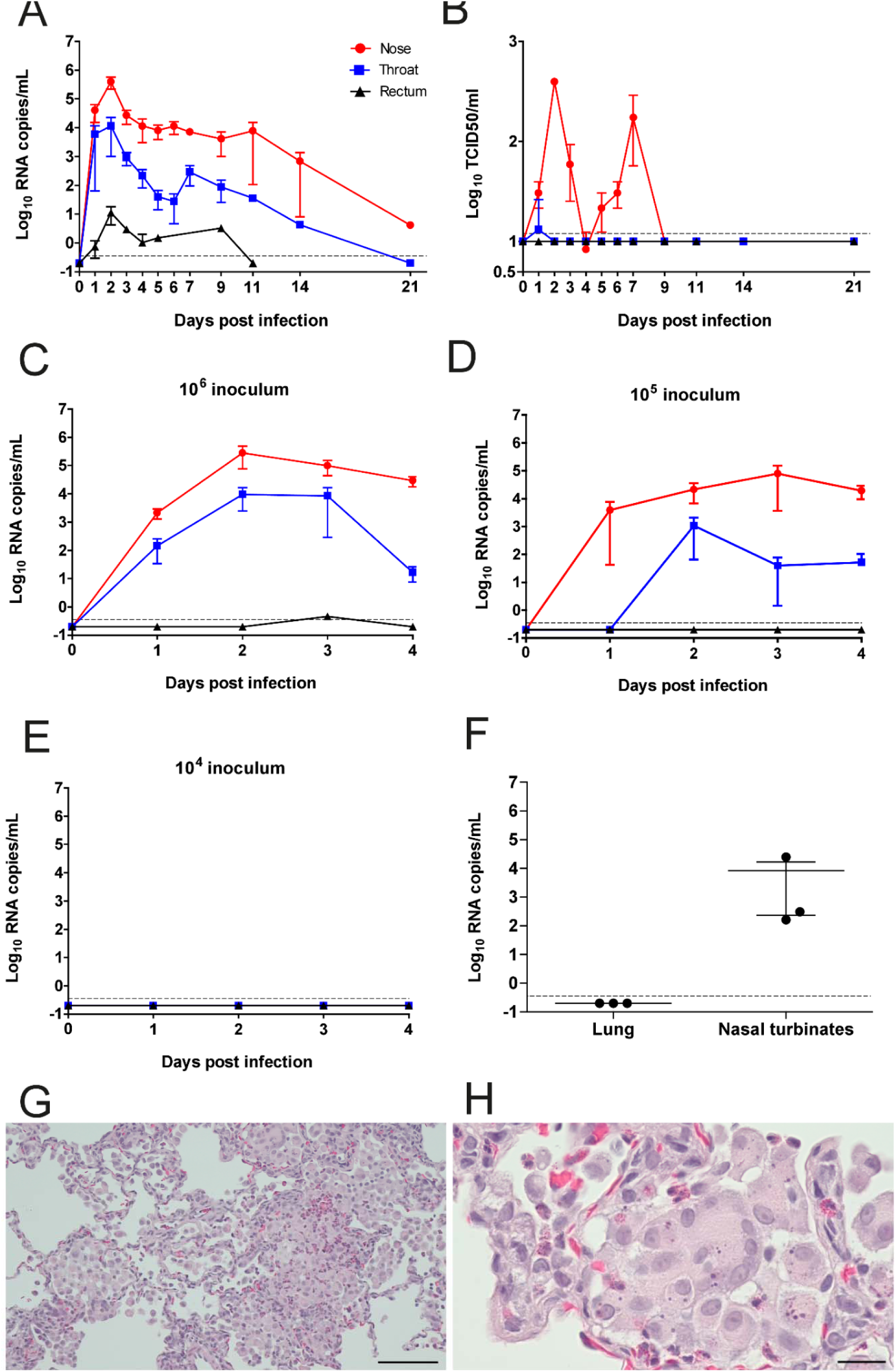
Susceptibility of rabbits to SARS-CoV-2 infection. Infection kinetics of (A) viral RNA and (B) authentic SARS-CoV-2 virus growth curves from rabbits inoculated with 10^6^ TCID_50_ and followed up fo r 21 days (C-E) Viral RNA growth curves in rabbits inoculated with either (C) 10^6^, (D) 10^5^, or (E) 10^4^ TCID_50_ and followed up for four days post infection. (F) Viral RNA in lung and nasal turbinates of 10^6^ TCID_50_ infected rabbits, sacrificed after four days. RNA detection limits were set at 3.5×10^−1^ RNA copies/ml, while live virus detection limit was 12 TCID_50_/ml. (G, H) Histopathological analysis of lungs from rabbits inoculated with 10^6^ TCID_50_, sacrificed after four days. (G) Alveolar thickening and inflammatory infiltrates. Scale indicates 100μm (H) Enlarged, syncytial cells in the alveolar lumina. Scale indicates 20μm.

Additionally, three groups of three animals were inoculated with either 10^4^, 10^5^ or 10^6^ TCID_50_ SARS-CoV-2 and swabs were taken for four days before the animals were euthanized and autopsied. All animals inoculated with 10^6^ TCID_50_ were viral RNA positive in the nose and throat for at least four days with a single animal positive in the rectum at day three (Figure 2, panel C). Animals inoculated with 10^5^ TCID_50_ were RNA positive in the nose for at least four days, for at least three days in the throat but not in the rectum (Figure 2, panel D). These animals also shed infectious virus in the nose for up to three days (mean shedding of 1.67 days, SD=1.53) and one animal shed virus two days post infection in the throat (data not shown). Animals inoculated with 10^4^ TCID_50_ did not shed any detectable viral RNA (Figure 2, panel E). Although nasal turbinates yielded on average 8.42×10^3^ RNA copies/ml, lung homogenates of animals inoculated with 10^6^ TCID_50_ virus were found viral RNA negative (Figure 2, panel F). Despite the fact that no viral RNA was detected in the lungs, histological examination of the lungs of infected animals sacrificed four days p.i. revealed a multifocal mild to moderate increase in alveolar macrophages in the alveolar lumina with multifocal presence of few neutrophils. Mainly associated with the terminal bronchioles, multifocal mild thickening of the septa, with infiltrations of neutrophils, eosinophils and occasional lymphocytes, plasma cells and macrophages was observed (Figure 2, panel G). Mild multifocal necrosis of alveolar epithelial cells and the presence of a few enlarged, syncytial cells in the alveolar lumina were seen (Figure 2, panel H). There was mild peribronchiolar and peribronchial lymphoplasmacytic infiltration with eosinophils and moderate to severe bronchus-associated lymphoid tissue proliferation (Supplementary figure 1, panel A). Some animals showed enlarged tracheo-bronchial lymph nodes consistent with mild lymphoid hyperplasia. In the nose, there was multifocal infiltration of moderate numbers of eosinophils and lymphoplasmacytic infiltrates in the olfactory epithelium (exocytosis) and in the lamina propria, alongside mild hyperplasia and hypertrophy of the olfactory epithelium (Supplementary figure 1, panel B). Mild eosinophilic exocytosis was present in the trachea.

## Conclusions

This study demonstrates that rabbits are susceptible to SARS-CoV-2. While the infection is asymptomatic, infectious virus with peak titers corresponding to ~10^3^ TCID_50_ could be detected up to day seven post inoculation in the nose. The minimum dose to establish productive infection was 10^5^ TCID_50_, indicating that virus transmission between rabbits may be less efficient compared to ferrets and hamsters. The use of young, immunocompetent, and healthy New Zealand White rabbits in this study however may not reflect virus shedding and disease in other rabbit breeds or rabbits at different ages. Thus, surveillance studies - including serological testing - may be needed to assess the presence of SARS-CoV-2 in farmed rabbits.

Viral shedding in rabbits occurred in a biphasic pattern, which was also observed for SARS-CoV-2 infected African green monkeys (*10*). This pattern is potentially linked to early innate immune responses that act within days, followed by adaptive responses that generally take one week to be activated. These observations are in line with recent findings that the presence of neutralizing serum antibodies in humans negatively correlates with infectious virus shedding, and that shedding of viral RNA outlasts shedding of infectious virus (*11*). The presence of eosinophils in the nose and lungs of infected animals suggests a possible helper T cell 2 (Th2)-mediated immune response. The preferential upper respiratory tract infection in the absence of robust replication in the lower respiratory tract of rabbits resembles what has been observed in experimentally inoculated ferrets (*5*).

The transmission of SARS-CoV-2 to mink caused viral spread between farm animals and spillover to humans, resulting in mass culling of mink to limit the spread of the virus (*6*). This study provides evidence of susceptibility of rabbits to SARS-CoV-2 infection warranting further investigations on the presence of SARS-CoV-2 in farmed rabbits.

## Acknowledgments

This research is partly financed by the Netherlands Organization for Health Research and Development (ZONMW) grant agreement 10150062010008 to B.L.H and co-funded by the PPP Allowance (grant agreement LSHM19136) made available by Health Holland, Top Sector Life Sciences & Health, to stimulate public-private partnerships.

## Author Bio

Anna Mykytyn is a PhD candidate at Erasmus Medical Centre, Rotterdam. Her research interests are the pathogenesis and transmission of coronaviruses including SARS-CoV-2.

## Supplementary

### Materials and Methods

#### Expression plasmids and cloning

Plasmids in pcDNA3.1 encoding human ACE2 (OHu20260), rabbit ACE2 (Clone ID OOb21562D), *Rhinolophus sinicus* ACE2 (ORh96277) and *Hipposideros armiger* ACE2 (Clone ID OHi02685) were ordered from GenScript. Codon-optimized cDNA encoding SARS-CoV-2 S glycoprotein (isolate Wuhan-Hu-1) with a C-terminal 19 amino acid deletion was synthesized and cloned into pCAGSS in between the EcoRI and BglII sites. pVSV-eGFP-dG (#31842), pMD2.G (#12259), pCAG-VSV-P (#64088), pCAG-VSV-L (#64085), pCAG-VSV-N (#64087) and pCAGGS-T7Opt (#65974) were ordered from Addgene. S expressing pCAGGS vectors were used for the production of pseudoviruses, as described below.

#### Cell lines

HEK-293T cells were maintained in Dulbecco’s Modified Eagle’s Medium (DMEM, Gibco) supplemented with 10% fetal bovine serum (FBS), 1X non-essential amino acids (Lonza), 1mM sodium pyruvate (Gibco), 2mM L-glutamine (Lonza), 100 μg/ml streptomycin (Lonza) and 100 U/ml penicillin. Cos-7, Vero, and VeroE6 cells were maintained in DMEM supplemented with 10% FBS, 1.5 mg/ml sodium bicarbonate (Lonza), 10mM HEPES (Lonza), 2mM L-glutamine, 100 μg/ml streptomycin and 100 U/ml penicillin. All cell lines were maintained at 37°C in a 5% CO_2_ humidified incubator.

#### VSV delta G rescue

The protocol for VSV-G pseudovirus rescue was adapted from Whelan and colleagues (*1*). Briefly, a 70% confluent 10 cm dish of HEK-293T cells was transfected with 10μg pVSV-eGFP- ΔG, 2μg pCAG-VSV-N (nucleocapsid), 2μg pCAG-VSV-L (polymerase), 2μg pMD2.G (glycoprotein, VSV-G), 2μg pCAG-VSV-P (phosphoprotein) and 2μg pCAGGS-T7Opt (T7 RNA polymerase) using polyethylenimine (PEI) at a ratio of 1:3 (DNA:PEI) in Opti-MEM I (1X) + GlutaMAX. Forty-eight hours post-transfection the supernatant was transferred onto new plates transfected 24 hours prior with VSV-G. After a further 48 hours, these plates were retransfected with VSV-G. After 24 hours the resulting pseudoviruses were collected, cleared by centrifugation at 2000 x g for 5 minutes, and stored at −80°C. Subsequently, VSV-G pseudovirus batches were produced by infecting VSV-G transfected HEK-293T cells with VSV-G pseudovirus at a MOI of 0.1. Titres were determined by preparing 10-fold serial dilutions in Opti-MEM I (1X) + GlutaMAX. Aliquots of each dilution were added to monolayers of 2 × 10^4^ Vero cells in the same medium in a 96-well plate. Three replicates were performed per pseudovirus stock. Plates were incubated at 37°C overnight and then scanned using an Amersham Typhoon scanner (GE Healthcare). Individual infected cells were quantified using ImageQuant TL software (GE Healthcare). All pseudovirus work was performed in a Class II Biosafety Cabinet under BSL-2 conditions at Erasmus Medical Center.

#### Coronavirus S pseudovirus production

For the production of SARS-CoV-2 S pseudovirus, HEK-293T cells were transfected with 10 μg S expression plasmids. Twenty-four hours post-transfection, the medium was replaced for OptiMEM I (1X) + GlutaMAX, and cells were infected at an MOI of 1 with VSV-G pseudovirus. Two hours post-infection, cells were washed three times with OptiMEM and replaced with medium containing anti-VSV-G neutralizing antibody (clone 8G5F11; Absolute Antibody) at a dilution of 1:50,000 to block remaining VSV-G pseudovirus. The supernatant was collected after 24 hours, cleared by centrifugation at 2000 x g for 5 minutes and stored at 4°C until use within 7 days. SARS-CoV-2 pseudovirus was titrated on VeroE6 cells as described above.

#### Virus stock

SARS-CoV-2 (isolate BetaCoV/Munich/BavPat1/2020; European Virus Archive Global #026V-03883; kindly provided by Dr. C. Drosten) was propagated on Vero E6 (ATCC® CRL 1586™) cells in OptiMEM I (1X) + GlutaMAX (Gibco), supplemented with penicillin (100 IU/mL) and streptomycin (100 IU/mL) at 37°C in a humidified CO2 incubator. Stocks were produced by infecting Vero E6 cells at a multiplicity of infection (MOI) of 0.01 and incubating the cells for 72 hours. The culture supernatant was cleared by centrifugation and stored in aliquots at −80°C. Stock titers were determined by titratin on VeroE6 cells. The TCID_50_ was calculated according to the method of Spearman & Kärber (ref?). All work with infectious SARS-CoV-2 was performed in a Class II Biosafety Cabinet under BSL-3 conditions at Erasmus Medical Center.

#### Pseudovirus and live virus infection *in vitro*

Cos-7 cells plated at 70% density in a 24 well format were transfected 24 hours after plating by dropwise addition of 500 ng ACE2 expression plasmids using a PEI ratio of 1:3 (DNA:PEI) in Opti-MEM I (1X) + GlutaMAX. After 24 hours, cells were washed twice and replaced with fresh Opti-MEM I (1X) + GlutaMAX prior to pseudovirus or live virus infection. Pseudovirus transduction was performed by infecting plates with 10^3^ VeroE6 titrated particles per well. Plates were incubated for 16 hours at 37°C before quantifying GFP-positive cells using an Amersham Typhoon scanner and ImageQuant TL software. Authentic virus infection was performed by adding 10^4^ TCID_50_ SARS-CoV-2 per well and incubating plates for 8 hours at 37°C. After incubation, cells were formalin fixed, permeabilized with 70% ethanol and stained with 1:1000 mouse anti-SARS nucleoprotein (Sino Biological) and 1:1000 rabbit anti-human ACE2 (Abcam), followed by 1:1000 goat anti-rabbit Alexa-Fluor 594, 1:1000 goat anti-mouse Alexa-Fluor 488, and 1:1000 TO-PRO3 (Thermo Fisher) to stain nuclei. Quantification of virus infected cells was performed using an Amersham Typhoon scanner and ImageQuant TL software as described above, while confocal imaging was performed on a LSM700 confocal microscope using ZEN software (Zeiss).

#### *In vivo* study design

Animal experiments were approved and performed according to the guidelines from the Institutional Animal Welfare Committee (AVD277002015283-WP04). The studies were performed under biosafety level 3 (BSL3) conditions. Three month-old New Zealand White rabbits (Oryctolagus cuniculus), specific pathogen free, and seronegative for SARS-CoV-2 were divided into four groups of three animals. Animals were inoculated under ketamine-medetomidine anesthesia intranasally with 1 ml SARS-CoV-2 (0.5 ml per nostril). Three groups were infected intranasally with respectively 10^4^, 10^5^ or 10^6^ TCID_50_ SARS-CoV-2 and monitored for four days post infection, and an additional three animals were infected with 10^6^ TCID_50_ SARS-CoV-2 and monitored for 21 days post infection. Animals monitored for four days were swabbed from the nose, throat, and rectum daily before being sacrificed for pathology on day 4 post infection. The remaining three 10^6^ TCID_50_ infected animals were followed until 21 days post infection with swabs taken on days zero to seven, then on days nine, 11, 14 and 21 post infection. In addition, on day 21 sera was collected from these animals for serological testing.

##### Plaque reduction neutralization test 50 (PRNT_50_)

Serum samples were tested for their neutralization capacity against SARS-CoV-2 as described before (*2*). Heat-inactivated (56 degrees Celsius for 30 minutes) samples were 2-fold serially diluted in Dulbecco modified Eagle medium supplemented with NaHCO3, HEPES buffer, penicillin, streptomycin, and 1% fetal bovine serum, starting at a dilution of 1:10 in 50 μL. Fifty μL virus suspension (~400 plaque-forming units) was added to each well and incubated at 37°C for 1 hour before transferring to VeroE6 cells. After incubation cells were washed and supplemented with medium, followed by an 8 hour incubation. After incubation, the cells were fixed with 4% formaldehyde/phosphate-buffered saline (PBS) and stained with polyclonal rabbit anti-SARS-CoV antibody (Sino Biological), followed by a secondary peroxidase-labeled goat anti-rabbit IgG (Dako). The signal was developed by using a precipitate forming 3,3′,5,5′ tetramethylbenzidine substrate (True Blue; Kirkegaard and Perry Laboratories) and the number of infected cells per well was quantified using an ImmunoSpot Image Analyzer (CTL Europe GmbH). The serum neutralization titer is the reciprocal of the highest dilution resulting in an infection reduction of >50% (PRNT_50_). A titer >20 was considered positive.

#### RNA extraction and qRT-PCR

Swabs, lung homogenates and nasal turbinate homogenates were thawed and centrifuged at 2,000 x g for 5 min. Sixty μL supernatant was lysed in 90 μL MagnaPure LC Lysis buffer (Roche) at room temperature for 10 minutes. RNA was extracted by incubating samples with 50 μL Agencourt AMPure XP beads (Beckman Coulter) for 15 minutes at room temperature, washing beads twice with 70% ethanol on a DynaMag-96 magnet (Invitrogen) and eluting in 30 μL ultrapure water. RNA copies per mL were determined by qRT-PCR using primers targeting the E gene (37) and compared to a counted RNA standard curve.

#### Histopathology

Alveolitis severity, bronchitis/bronchiolitis severity, tracheitis and rhinitis severity were scored: 0 = no inflammatory cells, 1 = few inflammatory cells, 2 = moderate number of inflammatory cells, 3 = many inflammatory cells. Alveolitis extent, 0 = 0%, 1 = <25%, 2 = 25-50%, 3 = >50%. Alveolar oedema presence, alveolar haemorrhage presence, type II pneumocyte hyperplasia presence, 0 = no, 1 = yes. Extent of peribronchial/perivascular cuffing, 0 = none, 1 = 1-2 cells thick, 2 = 3-10 cells thick, 3 = >10 cells thick.

#### Immunohistochemistry

Semiquantitative assessment of SARS-CoV-2 viral antigen expression in the lungs was performed as reported for SARS-CoV earlier *(3)* with a few amendments: for the alveoli, twenty-five arbitrarily chosen, 20x objective fields of lung parenchyma in each lung section were examined by light microscopy for the presence of SARS-CoV-2 nucleoprotein, without the knowledge of the allocation of the animals. The cumulative scores for each animal were presented as number of positive fields per 100 fields. For the bronchi and bronchioles, the percentage of positively staining bronchial and bronchiolar epithelium was estimated on every slide and the average of the four slides was taken to provide the score per animal. For the trachea and nose, the percentage of positively staining epithelium was estimated on every slide.

---

**Supplementary Figure 1.**
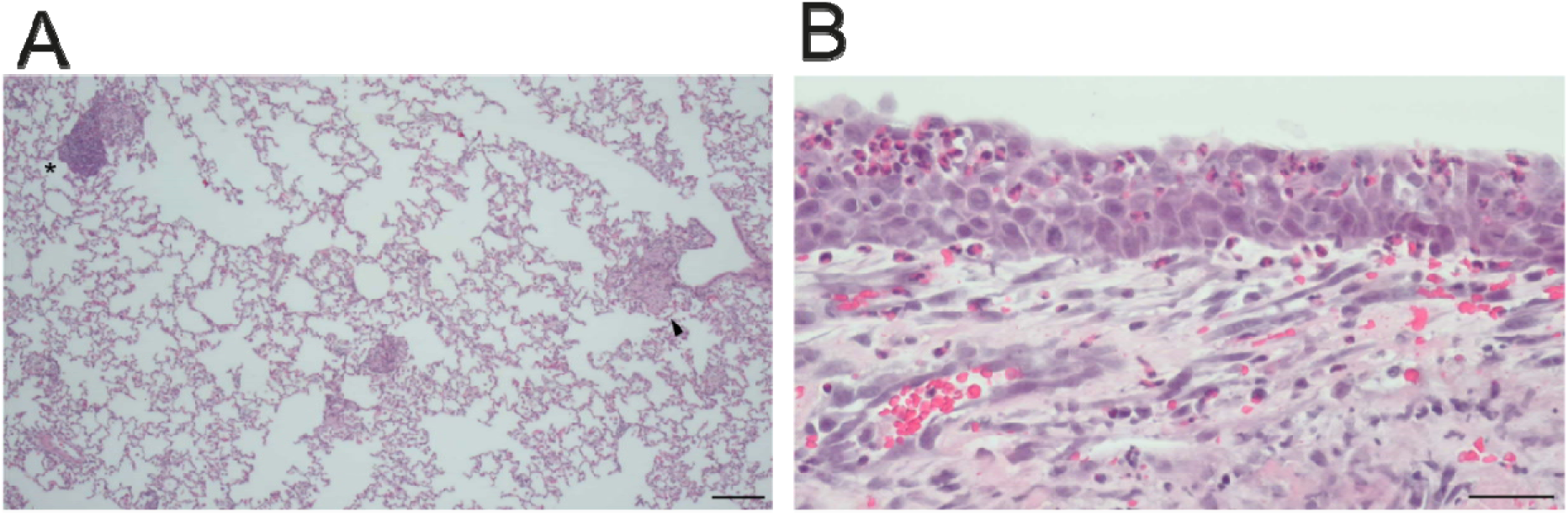
Lung histology of SARS-CoV-2 infected rabbits. (A) Lun pathology overview from rabbits inoculated with 10^6^ TCID_50_ and sacrificed 4 days post infection Arrow indicates thickening and asterisk bronchus-associated lymphoid tissue (BALT). Scale indicates 200μm. (B) Eosinophilic infiltrates in the nose. Scale indicates 40μm.

